# The consonance of chords: The effect of subsemitone changes to each of the notes in a trichord on its perceived consonance

**DOI:** 10.1101/2024.10.11.617763

**Authors:** J.M. Varkevisser, J.M. Kruitwagen, M. J. Spierings

## Abstract

Consonance plays a key role in how listeners experience music, but the factors influencing consonance perception remain a topic of discussion, for instance regarding the role of the familiarity of note combinations. Previous empirical studies of consonance have used common, familiar chords, making it harder to separate the effect of acoustic features from familiarity. Here we explored consonance perception in an unfamiliar, non-standardly tuned 0-5-10 trichord. A test was conducted in which 100 musicians and 100 nonmusicians compared the consonance of an original, non-standard, 0-5-10 chord and an adapted version, in which the lowest, middle or upper note was raised or lowered with 25 or 50 cents. According to a model based on continuous acoustic features, any adaptation to a 0-5-10 chord should yield a comparable decrease in consonance. We indeed found a general consonance decrease, but this was stronger for adaptations enlarging the distance between the lowest and the upper note, i.e. lowering the lowest or raising the highest note, than for other adaptations. These results deviate from the model’s predictions and from patterns observed in standard, more familiar chords. Future research should examine the generalizability of these findings to other chords than the 0-5-10 trichord.

## Introduction

Music is a human universal that plays a role in social and emotional contexts across populations (Mehr et al., 2019; Trehub et al., 2015). A key aspect of Western music, that partly drives the emotional response that it elicits, is the distinction between consonance and dissonance (Parncutt & Hair, 2011). Consonance describes sound combinations that listeners consider ‘pleasant’ and that are positively valenced, while dissonance describes sound combinations that listeners consider ‘unpleasant’ and that are negatively valenced (Parncutt & Hair, 2011). The concept of consonance and dissonance (C/D) can be applied to sounds perceived consecutively (melodic C/D) or simultaneously (simultaneous C/D), for instance in a chord.

In listening tasks, Western listeners show consistency in how they rate the simultaneous C/D of chords (Bowling et al., 2018; Parncutt et al., 2023). For instance, a C-E-G chord is perceived as highly consonant while a C-C#-D chord is perceived as highly dissonant. The factors that influence how a chord’s C/D is perceived have been the subject of ongoing discussion in the literature (Di Stefano et al., 2022; Harrison & Pearce, 2019). Several models have been proposed for predicting the perceived consonance of a chord based on its acoustic properties. These models typically highlight the acoustic features harmonicity and roughness as important contributors to C/D perception (Harrison & Pearce, 2019; Parncutt et al., 2023). Harmonicity here refers to the extent to which the frequencies of the notes in a chord resemble a harmonic spectrum, with the sound’s frequency components being integer multiples of the fundamental frequency (Harrison & Pearce, 2019; Parncutt, 1989) and roughness refers to the unpleasant sensation occurring when frequencies of tones are close together (Hutchinson & Knopoff, 1978).

To account for empirical data that deviate from the consonance predicted based on acoustic features, some models also incorporate familiarity or cultural factors, with more familiar combinations thought to be perceived as more consonant than unfamiliar sound combinations (Harrison & Pearce, 2019; Parncutt et al., 2023). However, other researchers argue that C/D perception of trichords can be fully explained by acoustic properties alone, if these properties also include the intervals between the notes in the chord (the number of semitones between the lower and middle note, and between the middle and upper note) and the ratio between these intervals (interval 1 : interval 2) (Cook & Fujisawa, 2006; Cook, 2017). In that case, according to this view, familiarity and cultural factors are unnecessary for explaining consonance ratings of chords.

These and other existing theories of C/D perception of trichords are, however, often based on, and empirically tested with, standardly-tuned chords that have integer semitone intervals. Examples include chords like C-E-G, with intervals of 4 and 3 semitones, or F-Ab-C, with intervals of 3 and 4 semitones, respectively. These types of chords are commonly used in music, which complicates disentangling possible effects of acoustic features and familiarity on the perceived consonance of these chords. In this study, we aim to empirically investigate trichord consonance perception using a non-standardly tuned trichord. Specifically, the notes in this trichord will not correspond to commonly used musical pitches (e.g., A4 at 440 Hz or G3 at 196 Hz), and the intervals between the notes will not be whole numbers of semitones. We believe that studying consonance in non-standardly tuned trichords with non-integer semitone intervals will provide valuable insights into how acoustic features influence consonance perception and we consider this study an initial effort in this direction.

In this study, we explore how small, subsemitone, changes to a non-standardly tuned 0-5-10 trichord affect its perceived consonance. Specifically, we alter the pitch of each of the notes in the trichord by raising or lowering them with 0.25 or 0.50 semitones (25 and 50 cents respectively; changes that should be well perceivable to both musicians and nonmusicians, Micheyl et al., 2006). These adjustments alter the harmonicity of the chord, and the (relative) interval sizes change. Since both the original and adapted versions of the chord are not commonly used in Western music, they are likely less influenced by people’s previous experiences with them. We examine how each of these pitch changes affects consonance perception relative to the original chord to find out whether changes to the lower, middle or upper note of a trichord have the largest impact on consonance perception. For the stimuli we use a 0-5-10 chord, which corresponds to a second inversion of a suspended fourth chord in music theory. We selected this chord because, unlike other chords, theoretical predictions suggest that both upward and downward shifts in the pitch of each of its notes should results in a comparable decrease in perceived consonance compared to the original trichord according to one of the models for the consonance of chords based on continuous acoustic features (Harrison & Pearce, 2018). If we find that not all shifts have an equally strong effect on consonance, this could give further insight into the acoustic properties shaping the perception of consonance in trichords.

We believe that three factors could lead to some shifts having a larger effect on consonance than others. We hypothesize that if (relative) interval sizes are key contributors to consonance perception, as proposed by Cook (2009; 2017), altering the middle note of the chord would have a larger impact on perceived consonance than other changes, as it modifies both intervals in the chord and leads to the most significant shift in the relationship between the two intervals compared to the original chord. Alternatively, changing the lowest note might have the largest effect, as the lowest note typically serves as the root of the chord, acting as a reference pitch for the other notes (Parncutt et al., 2023). However, the chord that was used in this experiment corresponds to the second inversion of a suspended chord, meaning that in this case the root of the chord is in the upper position. It is therefore also possible that changing the upper note of the chord, in this case, has a more pronounced effect on consonance perception than changes to the lowest or middle note.

## Methods

### Participants

We recruited 200 participants at the universities of Leiden and Delft and through personal connections of the researchers. 100 participants reported to be musicians (amateur or (semi)professional) and 100 reported to be nonmusicians. The mean age of the participants was 23.60 years (SD=4.57) and 54.5% of participants were women. All participants reported to have non-impaired hearing.

### Stimuli

All stimuli were based on a 0-5-10 chord, meaning the second note of the chord is five semitones higher than the lowest note and the third note is ten semitones higher. According to the Harrison and Pearce (2018) model, this chord has a stable decrease in consonance when any of the three notes is raised or lowered up to 0.50 semitones. Two sets of stimuli were made from the 0-5-10 chord with fundamental frequencies (F0) of 180 Hz and 190 Hz using Audacity (version 3.4.2). These frequencies differ from standard tuning notes but are closest to F#3 (185Hz). The frequencies used ensured that the chords had a relatively low pitch, making them pleasant for the participants to listen to (Di Stefano et al., 2022). Besides the original 0-5-10 chord, 12 adapted chords were made for both sets: in each adapted chord, one of the three notes was adapted in one of four ways: 0.50 semitones (50 cents) lower, 0.25 semitones (25 cents) lower, 0.25 semitones higher or 0.50 semitones higher. Each participant heard either both 0.25 semitone adaptations at 180 Hz and both 0.50 semitone adaptations at 190 Hz, or the other way around (both 0.25 semitone adaptations at 190 Hz and both 0.50 semitone adaptations at 180 Hz). Table 1 shows an overview of the stimuli.

**Table 1.**
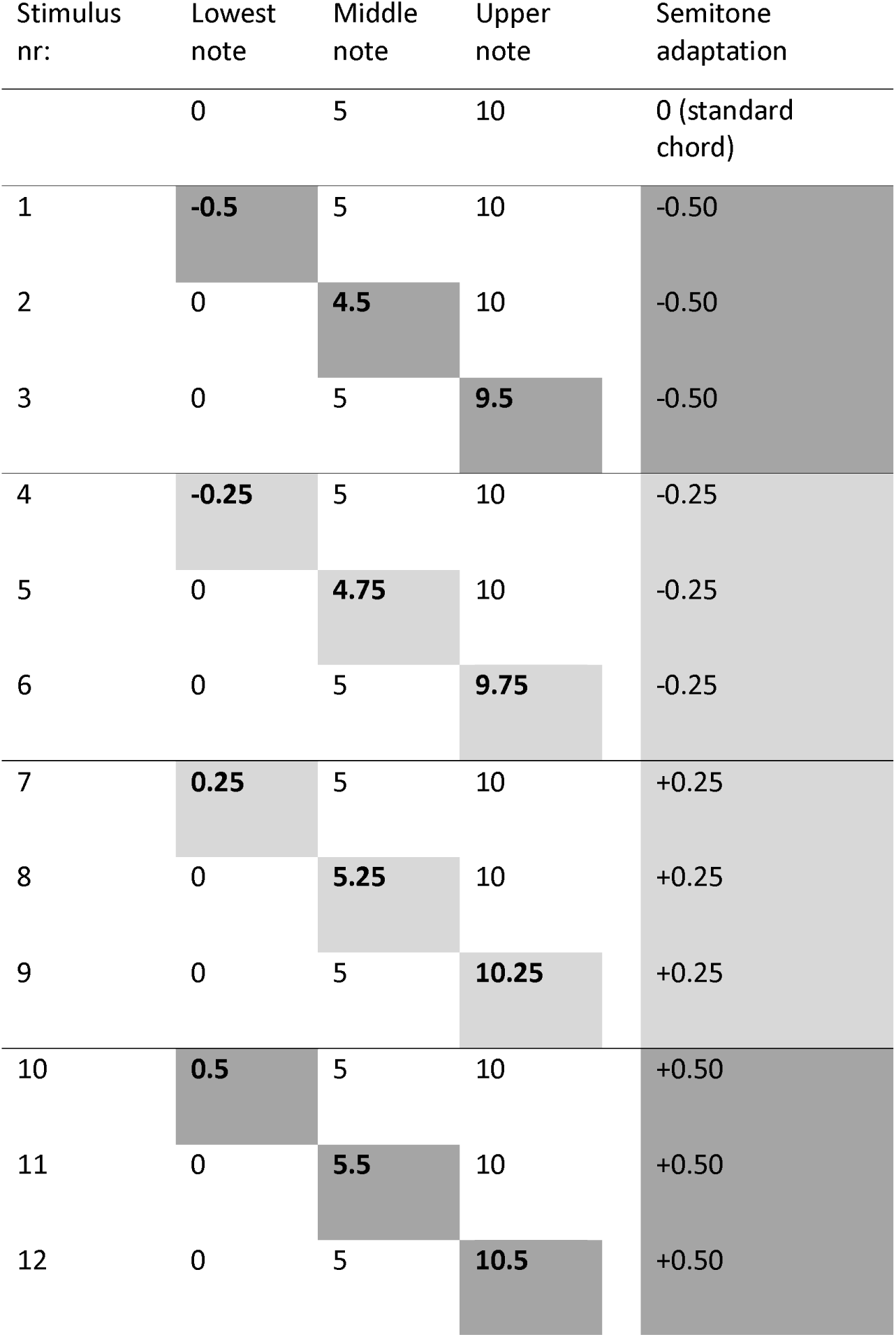
Overview of the stimuli used in the experiment. For each stimulus, the bold number corresponds to the note (lowest, middle or upper) that has been adapted relative to the original 0-5-10 chord. The two different shades of gray indicate how the stimuli belonging to the different stimulus sets (with F0 of 180 or 190 Hz) were distributed over participants: each participant heard the stimuli with the same color with the same fundamental frequency.

All stimuli were created using pre-recorded piano notes from the University of Iowa Electric Music Studios (https://theremin.music.uiowa.edu/MISpiano.html#). These notes have been recorded from a Steinway & Sons model B piano. The intermediate volume (annotated with ‘mf’) AIFF files of the Gb3, E3, and B4 were downloaded, as these are close to the frequencies needed for the stimuli. The sounds were adapted in Audacity so they had the required pitch and the three sounds were combined to create a trichord. The length of the sounds was adapted to create sounds of 1.6 seconds, with a fade-out where the mid-fade adjust was set at 70% to make the sounds resemble actual piano sounds. The stimuli were saved as WAV files.

Each trial in the experiment consisted of the presentation of the original 0-5-10 chord and one of the 12 adapted chords, both from the same stimulus set (so based on the same fundamental frequency). As previously mentioned, and as indicated in Table 1, there were two conditions regarding fundamental frequency that a participant could be assigned to. In Group A, the chords with the 0.25 semitone adaptations had a fundamental frequency of 180 Hz (light gray in table 1) and the chords with the 0.50 semitone adaptations had a fundamental frequency of 190 Hz (dark gray in table 1). For Group B, this was the other way around. To control for the effect of stimulus order in the experiment, twelve stimulus lists were made: six for Group A and six for Group B. The lists defined for each trial whether the original or the adapted chord was presented first, and in which order the adapted chords were presented. Every list contained each adapted chord three times for a total of 36 trials.

The lists were pseudorandomized to meet several conditions. First of all, since each adapted chord was presented three times, at least once the standard chord was presented first and at least once this adapted stimulus was presented first. Also, the first and last four stimuli of each list were counterbalanced between the twelve lists. This was done because it was anticipated that the first four and last four responses might not fully capture a participant’s true perception—either because they were just beginning (for the first four) or feeling fatigued (for the last four). Another restriction was that there could not be more than three consecutive stimuli from the same F0 group to reduce too much immediate repetition of the same original chord. Lastly, the three occurrences of a stimulus were always at least ten trials apart.

### Procedure

The experiment was executed in quiet places that were convenient for the participants. Acer Aspire 1 laptops were used to run the experiment, and stimuli were presented binaurally over Beyerdynamic DT 770 Pro headphones. Prior to the experiment, a sound level meter had been used to adjust the volume settings so that the stimuli were presented at 55 dB SPL. The experiment was designed and executed in E-Prime 3.0 (Psychology Software Tools, Pittsburgh, PA).

At the start of the experiment, we explained to the participants that they had to pay attention to the ‘harmonicity’ of complex sounds and explained ‘harmonicity’ as ‘how well the sounds within the chord go together’. We decided to use the term ‘harmonicity’ here because we thought this would be more intuitive for participants than ‘consonance’, and because in a previous study trichord ratings of this concept were highly correlated with ratings of ‘consonance’, but showed a higher inter-rater reliability across both musicians and non-musicians (Lahdelma & Eerola, 2020). As an example, sound files featuring a clear consonant (C4 and G4) and dissonant (C4 and Db4) interval were played. This was followed by four example trials (with intervals instead of chords) to familiarize the participants with the method of stimulus presentation and the response buttons.

A schematic overview of the experimental trials can be found in Figure 1. Each trial started with a fixation screen being displayed for 1000 milliseconds. Then, the first sound (the original or the adapted chord) played while the letter “A” was displayed on the screen. 3000 ms after the first sound started (thus ∼1.4 seconds after the first sound ended), the second sound (the original or the adapted chord) played while the letter “B” was displayed on the screen. From the start of the second sound, the participant had 7000 ms to indicate with the mouse which of these two chords they considered more ‘harmonious’, and to which extent, with the options 3, 2 and 1 on the left (sound A more harmonious than sound B), 0 (no difference between the two sounds in terms of harmonicity), and 1, 2 and 3 on the right (sound B more harmonious than sound A). If the participant failed to respond in time, the experiment continued to the next trial.

**Figure 1-.**
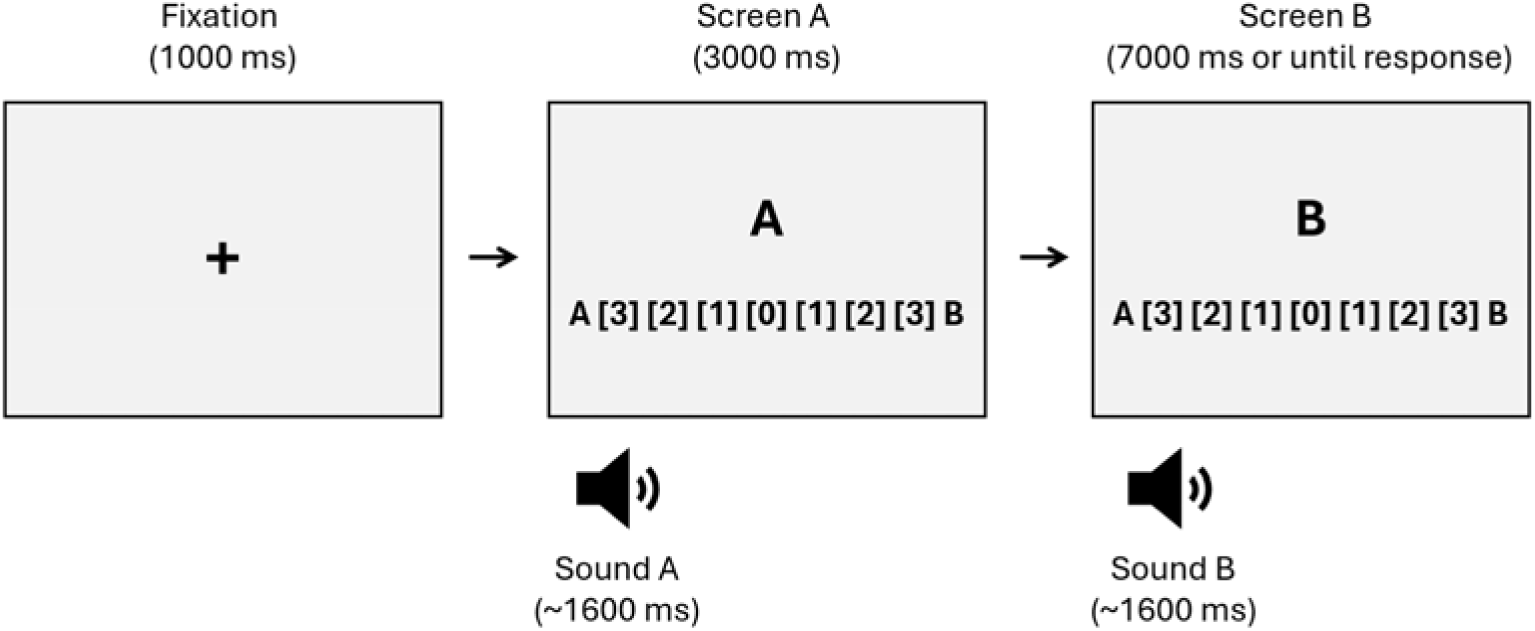
Schematic overview of one trial during the experiment. The content of the slides is simplified. The first slide displayed a fixation cross for 1000 milliseconds. The second slide showed the letter A and the response buttons while the first chord played. After 3000 milliseconds, the third slide appeared, displaying the letter B and the response buttons, which were now active. Simultaneously with the appearance of the third slide, the second chord was played.

During the example trials, the participant received visual feedback on whether they had responded in the correct manner (choosing one of the response options) or whether they were too slow with responding. For these example trials, the fixation time was 2000 ms so the participant had time to read the feedback. After the four example trials, the actual experiment started during which there was no feedback. Other than that, the example trials were exactly the same as the experimental trials. Halfway through the 36 trials, there was a break. The participant could decide when to continue to the second half of the trials by clicking the screen. After the experiment, the participant was asked several questions about their gender, age, musicianship and music preference within the E-Prime program. For the musicianship status, participants could choose between ‘nonmusician’, ‘music-loving nonmusician’, ‘amateur musician’, ‘serious amateur musician’, ‘semiprofessional musician’ or ‘professional musician’. This classification is based on the Ollen Musical Sophistication Index (Ollen, 2006) and has been shown to be an accurate single item measure of musical sophistication (Zhang & Schubert, 2019). The music preference genres that participants could choose from were ‘reflective and complex’, ‘intense and rebellious’, ‘upbeat and conventional’, and ‘energetic and rhythmic’, based on a factor analysis of music genres by Rentfrow and Gosling (2003). In total, the experiment took about eight minutes per participant after which they were rewarded with a snack.

### Statistical analysis

We analysed whether the adapated chords were perceived as more or less consonant than the original chords and whether the adaptations at the different note locations differentially affected consonance perception, by combining the button that the respondents chose and the chord that was presented first during a trial (the original or adapted chord) into a new variable, *PerceivedConsonanceDifference.* For this variable, a positive value indicates that the original chord was perceived as more consonant, a negative value indicates that the adapted chord was perceived as more consonant and a value of zero indicates that participants did not perceive the sounds as different in terms of consonance.

RStudio (R version 4.3.1, http://www.rstudio.com/) was used to model *PerceivedConsonanceDifference* with a cumulative link mixed model (CLMM) using the function “clmm” from package “ordinal” (Christensen, 2023). A CLMM is appropriate for ordinal logistic data such as the seven ordered categories of *PerceivedConsonanceDifference (−3, −2, −1, 0, 1, 2, 3)* and can deal with random effects. In the first basic model that was created, only the variables of interest *NoteLocation* (lowest, middle or upper note of the chord adapted) and *Adaptation* (min50, min25, plus25 or plus50 cents), and their interaction were included. After this basic model was created, other variables were added one at a time. These were *Gender* (male/female/other), *Age* (integer), *Musicianship* (Nonmusician/Musician, self-reported from six levels of musicianship), *Occurrence* (whether it was the first, second or third time a specific adapted chord was presented during the experiment), *OriginalChordFirst* (yes/no, whether the original chord was presented first within a trial), *AbsolutePitch* (yes/no, whether the participant does or does not have absolute pitch ability, self-reported), and *MusicPreference* (one of four meta-genres of music, self-reported). Each time a variable was added, this new model was compared to the previous model with an ANOVA (analysis of variance) and the variable was included if it significantly improved the model. In all models, *Subject* (participant number) and *ListName* (which stimulus list was presented, thus the order of stimulus presentation) were included as random effects. After the final model was created, the function “emmeans” from package “emmeans” (Lenth, 2023) was used for post hoc analyses while correcting for multiple comparisons with the Tukey method.

### Ethical note

Before the experiment, participants were asked to read and sign an informed consent form with information on the experiment and data handling, and on the participant’s option to stop with the experiment at any time. The experiment posed no known risks to the participants.

## Results

### General description of responses

Each of the 200 participants was presented with 36 trials, resulting in a total of 7200 trials. In 77 of these trials, participants did not answer within the allowed time of 7000 ms, leaving 7123 trials in the analysis. Table 2 shows the percentage of times participants responded with each of the response buttons during these trials. Table 3 shows these same responses, but now organised by whether they perceived the original chord as more consonant (positive categories) or the adapted chord as more consonant (negative categories), which represents the distribution on the *PerceivedConsonanceDifference* variable.

**Table 2.** Distribution of responses over response buttons. Participants had to select Button1, Button2 or Button3 if they thought the first sound was more consonant than the second sound they heard, with Button1 corresponding to a large difference in consonance and Button3 to a slight difference. Participants had to select Button4 if they did not hear a difference between the first and second sound in terms of consonance and Button5, Button6 or Button7 if they thought the second sound was more consonant than the first sound (with Button7 indicating a large difference).

| Button | Button1 | Button2 | Button3 | Button4 | Button5 | Button6 | Button7 |
| --- | --- | --- | --- | --- | --- | --- | --- |
| Number on<br>button | 3 | 2 | 1 | 0 | 1 | 2 | 3 |
| Percentage of<br>responses | 5.93% | 13.72% | 18.74% | 29.36% | 18.32% | 9.61% | 3.25% |

**Table 3.** Distribution of responses on the categories of the *PerceivedConsonanceDifference* variable. A positive response indicates that the original chord was perceived as more consonant (61.78% of responses), a negative response indicates that the adapted chord was perceived as more consonant (7.79%), and a “0” response indicates that the original chord and adapted chord were perceived as being equally consonant (29.36%).

| Category | -3 | -2 | -1 | 0 | 1 | 2 | 3 |
| --- | --- | --- | --- | --- | --- | --- | --- |
| Percentage of responses | 0.36% | 1.53% | 5.90% | 29.36% | 31.15% | 21.81% | 8.82% |

### Effect of adaptations at different note locations on consonance perception

The final model for the relationship between consonance perception (*PerceivedConsonanceDifference)* and the adaptations at the different note locations (*Adaptation, NoteLocation* and their interaction) also included the variables *Musicianship*, *Gender*, *OriginalChordFirst* and *Occurrence* (Table 4). Not included were *Age*, *AbsolutePitch* and *MusicPreference*.

**Table 4.**
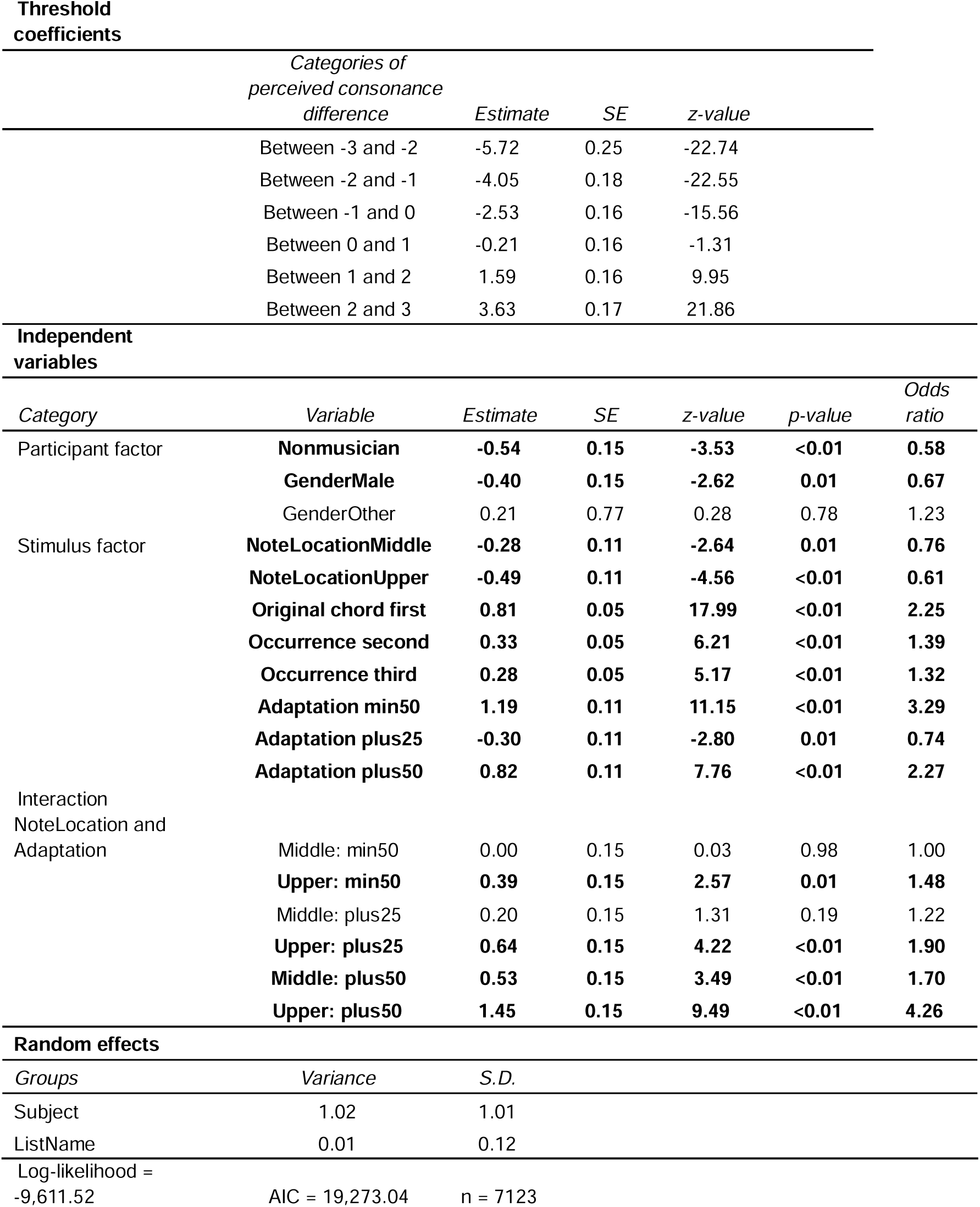
Results of the cumulative link mixed model (CLMM) for the effect of adaptations at different note locations in a trichord on perceived differences in consonance. The odds ratio (OR) reflects the association between the predictor variable and the likelihood of the response falling into a higher category versus a lower one. For example, when the original chord is presented first, the odds are 2.25x higher that the score on *PerceivedConsonanceDifference* would be in a higher category than when the original chord is presented second. Significant effects are marked in bold (p < 0.05).

There were significant main effects of *NoteLocation* and *Adaptation.* If the middle or upper note was changed in the adapted chord, the odds were lower that a participant would give a higher response than when the lowest note was changed (Table 3, middle: OR=0.76, estimate=-0.28, SE=0.11, p<0.01, upper: OR=0.61, estimate=-0.49, SE=0.11, p<0.001). Compared to min25, adaptations of min50 and plus50 increased the odds that a higher response was given (min50: OR=3.29, estimate=1.19, SE=0.11, p<0.001, plus50: OR=2.27, estimate=0.82, SE=0.11, p<0.001) and an adaptation of plus25 decreased the odds that a higher response was given (OR=0.74, estimate=-0.30, SE=0.11, p<0.01). There was, however, a significant interaction between adaptation and note location, indicating that the effect of the different adaptations depended on the note that was adapted. To further explore this interaction, we conducted post hoc analyses to address two questions: first whether the same adaptation had different effects when applied to the upper, middle, or lower note of the trichord and second, whether there was a difference in effect if the same note was either raised or lowered.

First, the post hoc analysis showed that the effect of the min25 and plus50 adaptations significantly varied between note locations (Figure 2, Table 5). When the lowest note was changed 25 cents down, responses were significantly higher than when the upper note was changed 25 cents down (estimate=0.49, SE=0.11, p<0.001). Changing the upper note 50 cents up resulted in significantly higher responses compared to changing either the lowest note 50 cents up (estimate=-0.96, SE=0.11, p<0.001) or the middle note 50 cents up (estimate=-2.17, p<0.001).

**Figure 2-.**
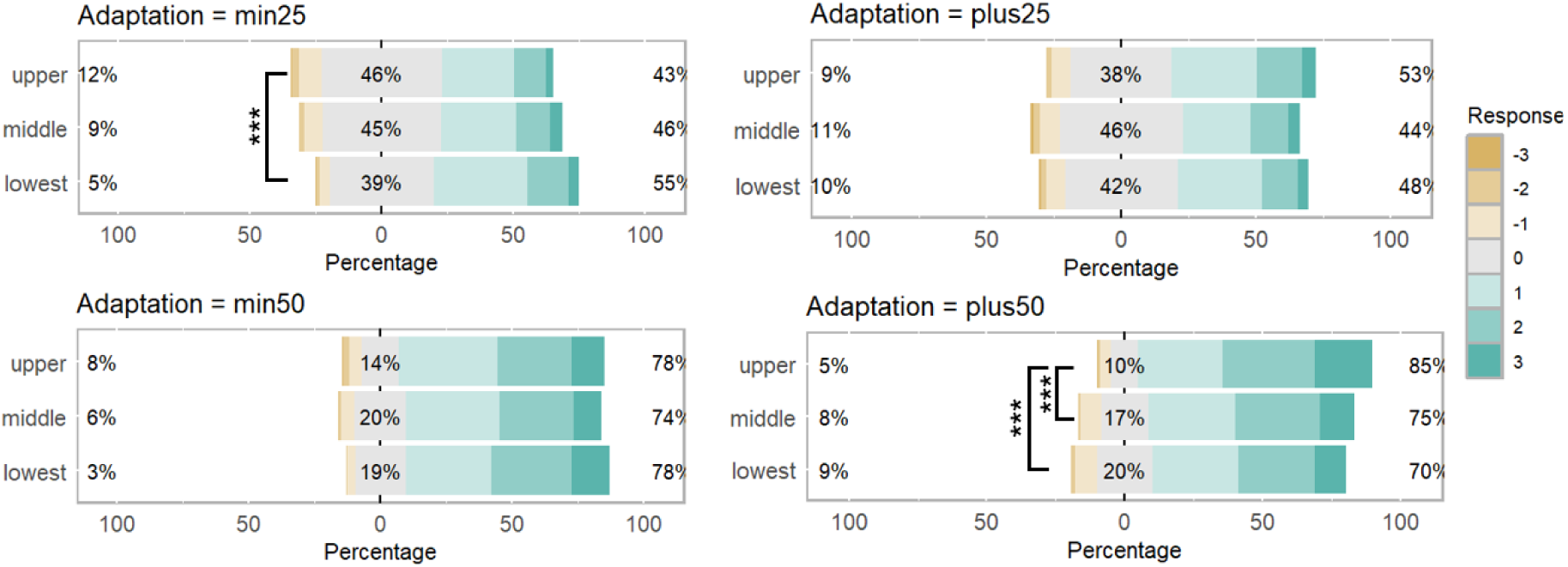
Comparisons of the effect of each adaptation per note location on perceived consonance difference. Percentages of responses on the *PerceivedConsonanceDifference* variable given for each adaptation (min25, min50, plus25, plus50) per note location (upper, middle, and lowest). The percentages are centered around the answer “0”. For each bar, three percentages are shown. The percentage on the left is the percentage of answers where the adapted chord was perceived as more consonant than the original chord (responses that were −3, −2 or −1). The percentage in the gray part of the bar is the percentage of “0” answers. The percentage on the right is the percentage of answers where the original chord was perceived as more consonant than the adapted chord (1, 2, or 3). ***: p<0.001

**Table 5.** Post hoc comparison of the effect of each adaptation per note location on the perceived consonance difference. *C*omparisons for CLMM of perceived consonance difference (Table 4). P-value adjustment for multiple comparisons with Tukey method. Significant effects (p<0.05) are marked in bold.

| *min50* | estimate | SE | z-value | p-value |
| --- | --- | --- | --- | --- |
| lowest min50 - middle min50 | 0.28 | 0.11 | 2.6 | 0.28 |
| lowest min50 - upper min50 | 0.10 | 0.11 | 0.89 | 1.00 |
| middle min50 - upper min50 | -0.18 | 0.11 | -1.69 | 0.87 |
| *min25* |  |  |  |  |
| lowest min25 - middle min25 | 0.28 | 0.11 | 2.64 | 0.26 |
| **lowest min25 - upper min25** | **0.49** | **0.11** | **4.56** | **<0.001** |
| middle min25 - upper min25 | 0.21 | 0.11 | 1.89 | 0.76 |
| *plus25* |  |  |  |  |
| lowest plus25 - middle plus25 | 0.08 | 0.11 | 0.78 | 1.00 |
| lowest plus25 - upper plus25 | -0.15 | 0.11 | -1.41 | 0.96 |
| middle plus25 - upper plus25 | -0.23 | 0.11 | -2.18 | 0.56 |
| *plus50* |  |  |  |  |
| lowest plus50 - middle plus50 | -0.25 | 0.11 | -2.3 | 0.48 |
| **lowest plus50 - upper plus50** | **-0.96** | **0.11** | **-8.84** | **<0.001** |
| **middle plus50 - upper plus50** | **-0.71** | **0.11** | **-6.58** | **<0.001** |

Second, the post hoc analysis showed that responses were significantly higher when the upper note was changed 50 cents up rather than 50 cents down (estimate=-0.69, SE=0.11, p<0.001), while the responses were significantly lower when the lowest note was changed 50 cents up rather than 50 cents down (estimate=0.37, SE=0.11, p=0.03, Figure 3, Table 6).

**Figure 3-.**
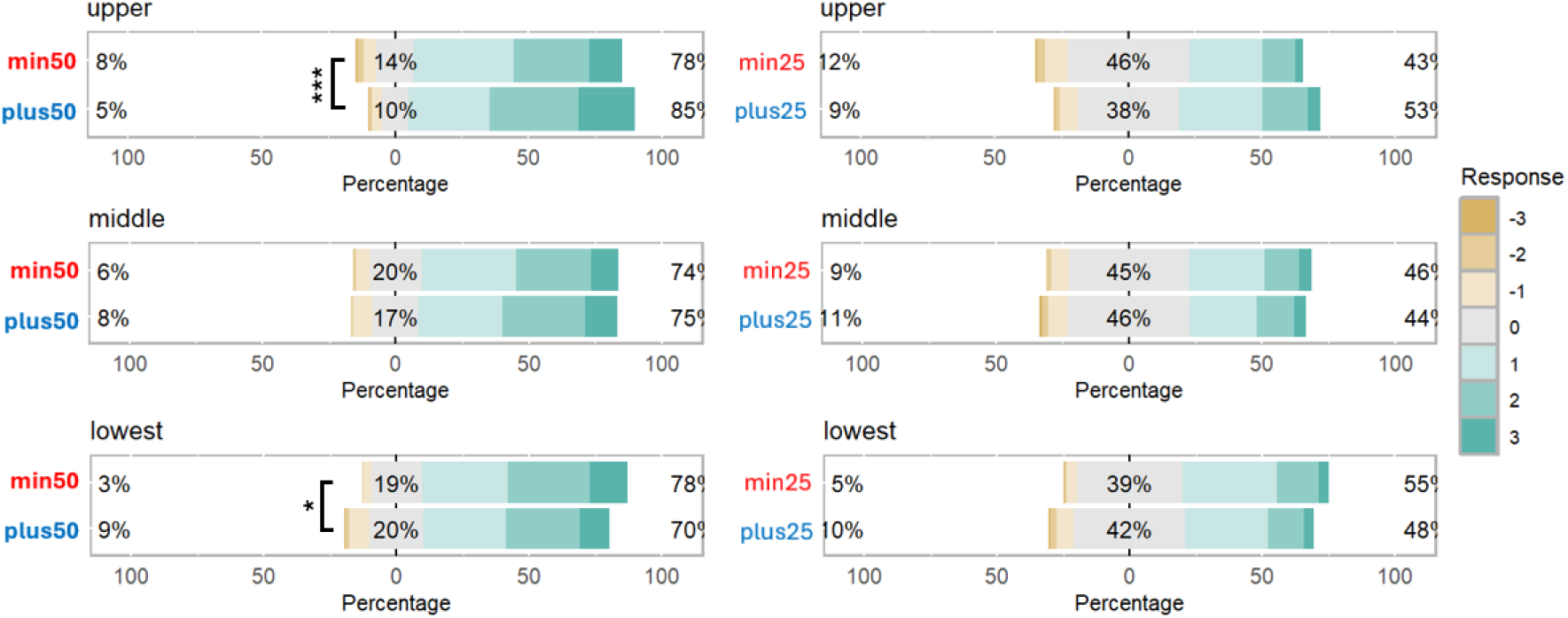
Comparisons between the adaptations upward and downward per note location. Percentages of responses on the *PerceivedConsonanceDifference* variable for the adaptations upward (plus25, plus50) and downward (min25, min50) per note location (upper, middle and lowest). For each note location, the effect of the direction of the adaptation is visualized. *: p<0.05, ***: p<0.001.

**Table 6.**
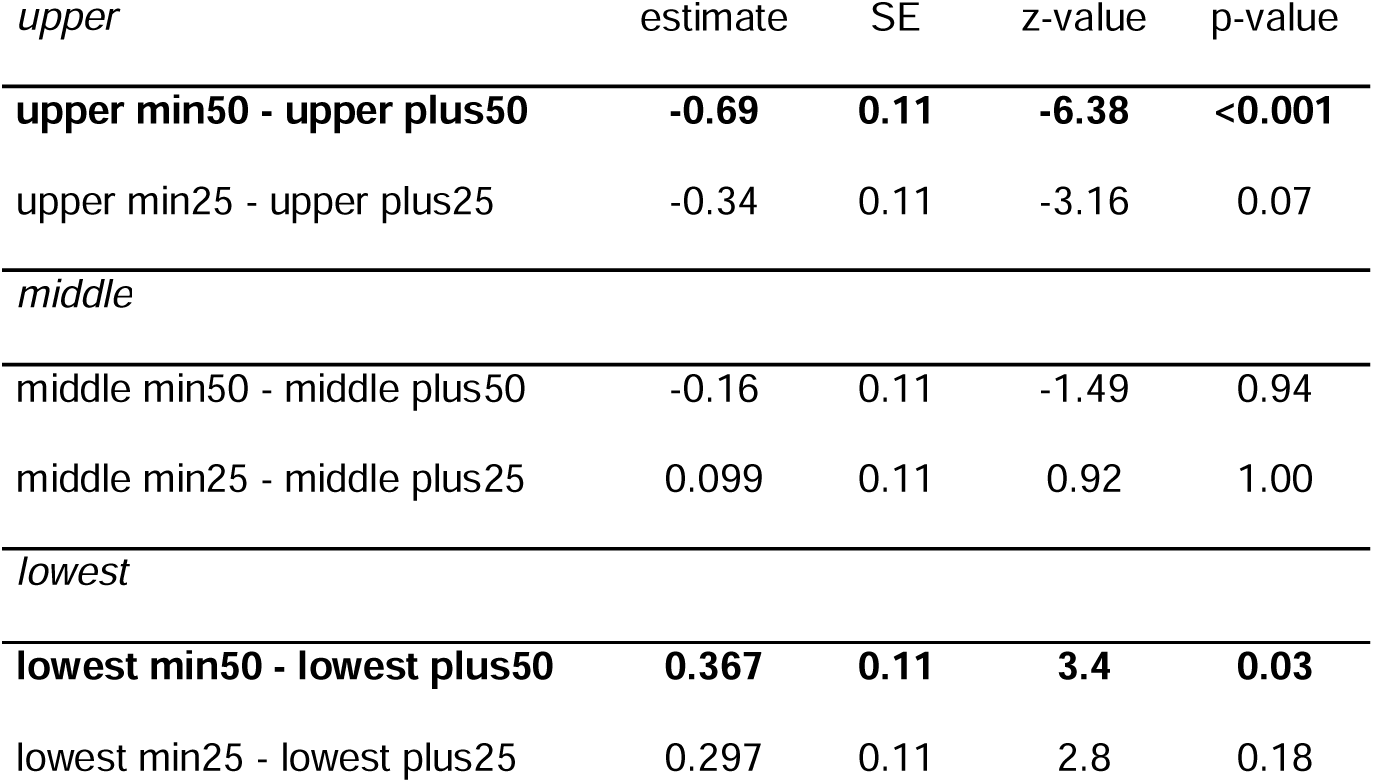
Post hoc comparison of the effect of the direction of the adaptations (upwards or downwards) per note location. Comparisons for CLMM of perceived consonance difference (see Table 4). P-value adjustment for multiple comparisons with Tukey method. Significant interaction effects (p<0.05) are marked in bold.

| *upper* | estimate | SE | z-value | p-value |
| --- | --- | --- | --- | --- |
| **upper min50 - upper plus50** | **-0.69** | **0.11** | **-6.38** | **<0.001** |
| upper min25 - upper plus25 | -0.34 | 0.11 | -3.16 | 0.07 |
| *middle* |  |  |  |  |
| middle min50 - middle plus50 | -0.16 | 0.11 | -1.49 | 0.94 |
| middle min25 - middle plus25 | 0.099 | 0.11 | 0.92 | 1.00 |
| *lowest* |  |  |  |  |
| **lowest min50 - lowest plus50** | **0.367** | **0.11** | **3.4** | **0.03** |
| lowest min25 - lowest plus25 | 0.297 | 0.11 | 2.8 | 0.18 |

### Additional significant effects

According to the CLMM, two participant factors had a significant effect on the perceived consonance difference: gender and musicianship (Table 3). Men had lower odds of selecting higher response categories compared to women (OR=0.67, estimate=-0.40, SE=0.15, p<0.01) and nonmusicians had lower odds compared to musicians (OR=0.58, estimate=-0.54, SE=0.15, p<0.001). Besides these participant factors, the odds of selecting a higher category increased with the second and third occurrence of the chord compared to the first occurrence (second occurrence: OR=1.39, estimate=0.33, SE=0.05, p<0.001, third occurrence: OR=1.32, estimate=0.28, SE=0.05, p<0.001), and were higher if the original chord was presented first rather than the adapted chord (OR=2.25, estimate=0.81, SE=0.05, p<0.001, Table 3).

## Discussion

The aim of this study was to investigate how subsemitone upward and downward adaptations to each of the notes in a non-standardly tuned 0-5-10 trichord affect its perceived consonance, compared to the original, non-adapted 0-5-10 chord. Our results indicate that in most cases participants either perceived the adapted trichords as less consonant than the original trichord (approximately 62% of the trials), or they did not perceive a difference in consonance (approx. 29%), while only in few cases (approx. 8%) they perceived the adapted trichord as more consonant. This aligns with the trichord consonance model proposed by Harrison and Pearce (2018), which predicts that small changes to each of the notes of a 0-5-10 trichord reduce its perceived consonance compared to the original trichord. However, contrary to this model’s prediction, not all of the up- and downward adaptations resulted in a comparable decrease in consonance.

We hypothesized that if the different adaptations would not lead to a comparable decrease in consonance, changes to the middle note would have the largest effect on consonance perception due to its influence on the (relative) interval sizes within the chord. Alternatively, we considered that the lowest or upper note might have a larger effect than the other notes, given that the lower note typically serves as the root in root position chords, and the upper note functions as the root in second inversions, such as the 0-5-10 chord used in this experiment. However, contrary to our hypotheses, we found that the effect of the adaptations depended on both the position of the note that was altered and the direction of the adaptation. Participants reported a larger consonance decrease from the original to the adapted chord when the lowest note was lowered compared to when it was raised, and when the lowest note was lowered compared to when the upper note was lowered. Furthermore, raising the upper note led to a larger consonance decrease than lowering it, and raising the upper note caused a more pronounced consonance decrease than raising either the lowest or middle note.

The general decrease in consonance found with any of the changes to the 0-5-10 chord may be related to the chord’s structure, which consists of two equidistant intervals of five semitones each. Even spacing of harmonics, such as in equidistant intervals between notes, reduces interference and enhances consonance (Harrison & Pearce, 2019). The changes applied to the chord in this experiment disrupted the evenness, likely thereby reducing its consonance. The most pronounced decrease in perceived consonance was found when the lowest note was lowered or the highest note of the trichord was raised. Generally, larger intervals are thought to produce less roughness than smaller intervals (Parncutt et al., 2023), which would contrast with our findings, but this is only found in smaller intervals (of 20 to 200 Hz, De Baene et al., 2004) than the ones used in our study. Moreover, enlarging intervals by changes to the middle note did not have the same effect as changes to the lowest or upper note, showing that consonance perception in our experiment was not mainly driven by the size of the intervals between two adjacent notes. Finally, our results do not match the observation in standard chords that chords with a lower interval larger than the upper interval are typically more positively valenced than chords with a smaller interval below the upper interval (Cook, 2009; Parncutt et al., 2023). In contrast, we observed a stronger decrease in consonance when the lowest note was lowered – creating a larger lower than upper interval - than when it was raised, which resulted in a larger upper than lower interval, and whether the middle note was raised or lowered did not lead to significant differences in the perceived consonance. This suggests that the decrease in consonance in our study was mainly driven by the increased distance between the upper and lowest note of the trichord.

Our results demonstrate that although a model based on acoustic features (Harrison & Pearce, 2018) predicts a comparable decrease in consonance with any change to a note in a 0-5-10 trichord, additional patterns arise when this is tested empirically. Cultural factors or learning effects are sometimes referred to in other cases where empirical data diverge from model predictions. In this case, however, cultural influences seem less relevant, due to the use of a non-standardly tuned chord and non-integer semitone intervals, which are uncommon in Western music. It is possible, though, that some of the adaptations used in this experiment caused the stimulus chord to resemble a familiar chord more closely than others, suggesting that familiarity might still play a role in our results.

Two participant-related factors significantly influenced the perceived consonance difference. First of all, musicians were more likely to report a larger consonance difference than nonmusicians, which could be due to their enhanced ability to detect small pitch deviations (Micheyl et al., 2006). Additionally, women reported larger differences in consonance than men, which is in line with previous studies that found a larger response to harmonics in sound in women than in men (Krizman et al., 2021). On the other hand, no significant effect of music preference was found, while it could be theorized that participants who favour the meta-genre “Intense and Rebellious (for example Rock, Alternative, Heavy metal)” have more exposure to dissonance-rich music, which could result in a different experience of consonance (Harrison & Pearce, 2019). However, many of our participants reported difficulty with selecting one meta-genre of music, as they engaged with multiple music styles. No effect of age was found among our participants, though this was expected as only 6% of participants were over 30. Similarly, no effect of absolute pitch was observed, with only 5% of participants reporting having this ability.

The perceived consonance difference was also influenced by two stimulus factors. When the original chord was presented first, the perceived consonance difference was larger compared to when it was presented second. This may be related to the concept of resolution, which occurs when an unpleasant-sounding chord is followed by a pleasant-sounding one (Parncutt & Hair, 2011). This pleasant sensation might have led participants to report a relatively smaller consonance difference between the two chords. Additionally, each chord was presented three times during the experiment and during later occurrences, the perceived consonance differences were larger. It is possible that as participants became more familiar with the small adaptations throughout the experiment, their ability to discriminate between the chords improved.

Further research is recommended to study the generalizability of this study’s findings. The current study only included one chord and was conducted exclusively with Western listeners. Therefore, it remains uncertain whether the effects that we found are specific to the 0-5-10 chord or would also be found with other chords. To draw more general conclusions, further experiment should include a variety of chords, while taking into account the variable theoretical consonance landscape of these chords. Additionally, comparable future studies should also include non-Western participants that are less familiar with the Western tonal framework to draw more general conclusions about human consonance perception.

In conclusion, our study demonstrates that subsemitone changes to each of the notes of a non-standardly tuned 0-5-10 trichord generally result in a decrease in perceived consonance compared to the original, unaltered chord. The largest decrease in consonance was found for changes that increased the distance between the lowest and upper note of the trichord. Other patterns that have been observed in studies with standard, more familiar chords, e.g. related to the effect of the relative interval size of the notes, were not observed in our study. Future studies should further explore the factors underlying the patterns found in this study, for instance by investigating whether these patterns are specific to the 0-5-10 chord used in this study or apply more generally to other chords as well. Eventually these insights could help refine existing models for the perceived consonance of trichords.

